# Substantial near-infrared radiation-driven photosynthesis of chlorophyll *f*-containing cyanobacteria in a natural habitat

**DOI:** 10.1101/750174

**Authors:** Michael Kühl, Erik Trampe, Maria Mosshammer, Michael Johnson, Anthony W. D. Larkum, Klaus Koren

**Affiliations:** Marine Biological Section, Department of Biology, University of Copenhagen, Denmark; Climate Change Cluster, University of Technology Sydney, Ultimo, Australia; iThree Institute, University of Technology Sydney, Australia; Centre for Water Technology, Section for Microbiology, Department of Bioscience, University of Aarhus, Denmark Aarhus C, Denmark

## Abstract

Far-red absorbing chlorophylls are constitutively present as Chl *d* in the cyanobacterium *Acaryochloris marina*, or dynamically expressed by synthesis of Chl *f* and red-shifted phycobilins via far-red light photoacclimation in a range of cyanobacteria, which enables them to use near-infrared-radiation (NIR) for oxygenic photosynthesis. While the biochemistry and molecular physiology of Chl *f*-containing cyanobacteria has been unraveled in culture studies, their ecological significance remains unexplored and no data on their *in situ* activity exist. With a novel combination of hyperspectral imaging, confocal laser scanning microscopy, and nanoparticle-based O_2_ imaging, we demonstrate substantial NIR-driven oxygenic photosynthesis by endolithic, Chl *f*-containing cyanobacteria within natural beachrock biofilms that are widespread on (sub)tropical coastlines. This indicates an important role of NIR-driven oxygenic photosynthesis in primary production of endolithic and other shaded habitats.

**Impact statement:** Cyanobacteria with chlorophyll *f* show substantial near-infrared radiation-driven photosynthesis in intertidal habitats.

## Introduction

The persisting textbook notion that oxygenic photosynthesis is mainly driven by visible wavelengths of light (400-700 nm) and chlorophyll (Chl) *a* as the major photopigment is challenged. Recent findings indicate that cyanobacteria with red-shifted chlorophylls and phycobilins capable of harvesting near infrared radiation (NIR) at wavelengths >700-760 nm and exhibiting a pronounced plasticity in their photoacclimatory responses (*Gan et al., 2014*; *Gan and Bryant, 2015*) are widespread in natural habitats (*Gan et al., 2015*; *Zhang et al., 2019*; *Behrendt et al., 2019*). Besides the Chl *d*-containing cyanobacterium *Acaryochloris marina*, which was originally isolated from tropical ascidians (*Miyashita et al., 2014*) but has now been found in many other habitats (*Behrendt et al., 2011*; *Zhang et al., 2019*), the discovery of Chl *f* (*Chen et al., 2010*) and its occurrence in many different cyanobacteria (*Gan et al., 2015*) has triggered a substantial amount of research on the biochemistry and molecular physiology of Chl *f*-containing cyanobacterial strains (*Airs et al., 2014*; *Allakhverdiev et al., 2016*; *Chen, 2014*; *Ho et al., 2016*; *Nürnberg et al., 2018*). In comparison, the *in situ* distribution and activity of Chl *f*-containing cyanobacteria and their role in primary productivity remain largely unexplored. Here we used a novel combination of hyperspectral imaging, confocal laser scanning microscopy, and chemical imaging of O_2_ for high-resolution mapping of the distribution of Chl *f*–containing cyanobacteria and their NIR-driven oxygenic photosynthesis in an intertidal beachrock habitat. Our study gives novel insight into the ecological niche and importance of endolithic Chl *f*-containing cyanobacteria, and indicates that high rates of NIR-driven oxygenic photosynthesis can contribute to primary production in natural biofilm habitats.

With an *in vivo* absorption range of 700-760 nm, Chl *f* is the most red-shifted chlorophyll, which was first found in the filamentous cyanobacterium *Halomicronema hongdechloris* isolated from stromatolites in Western Australia (*Chen et al., 2010*; *Chen et al. 2013*), and in an unicellular cyanobacterium (strain KC1 related to *Aphanocapsa* spp.) isolated from Lake Biwa, Japan (*Akutsu et al., 2011*). However, Bryant and coworkers discovered that the ability to synthesize Chl *f*, far-red shifted phycobilins, and small amounts of Chl *d* can be induced in many different cyanobacteria, including representatives from all 5 major subdivisons, when grown under far-red light-enriched conditions (*Gan et al., 2014*; *Gan et al. 2015*). Such far-red light photoacclimation (FaRLiP) involves remodeling of the photosynthetic apparatus via synthesis and modification of pigments and pigment-protein complexes. This remodeling is primarily regulated at the transcriptional level via upregulation of paralogous photosynthesis-related genes in a 21-gene cluster, which seems largely conserved in cyanobacteria exhibiting FaRLiP (*Gan and Bryant, 2015*) and contains genes for a red phytochrome-triggered control cascade of FaRLiP (*Zhao et al., 2015*). Recently, it was shown that FaRLiP also involves the modification and inclusion of Chl *f* in both PSI and PSII reaction centers (*Itoh et al., 2015*; *Nürnberg et al., 2018*).

There is a growing database of cyanobacteria and habitats wherein Chl *f* has been detected (Table S1). However, only three studies have reported on the actual distribution and niche of Chl *f*-containing cyanobacteria in their natural habitats (*Trampe and Kühl 2016*; *Ohkubo and Miyashita, 2017*; *Behrendt et al., 2019*), and hitherto NIR-driven oxygenic photosynthesis by Chl *f*-containing cyanobacteria has not been demonstrated *in situ*. This is experimentally challenging, as cyanobacteria with FaRLiP respond to the local light microenvironment and typically occur in dense proximity with other oxygenic phototrophs harboring a plethora of photopigments fueling Chl *a*-based oxygenic photosynthesis by visible light (400-700 nm) (*Okhubo and Miyashita, 2017*; *Behrendt et al., 2015*). Consequently, photosynthetic activity of Chl *f*-containing cyanobacteria has only been measured on strains (e.g. *Gan et al., 2014*; *Nürnberg et al., 2018*) or enrichments isolated from their natural habitat (e.g. *Behrendt et al., 2015*). Detailed microscopic investigation of pigmentation in cyanobacteria with Chl *f* has also largely been limited to culture material (*Majumder et al., 2017*; *Zhang et al., 2019*).

Beachrock is a widespread sedimentary rock formation on (sub)tropical, intertidal shorelines, where a mixture of biogeochemical processes cement carbonate sands together into a porous solid matrix (*Vousdoukas et al., 2007*). The upper surface of light-exposed beachrock is colonized by dense microbial biofilms dominated by cyanobacteria (*Cribb, 1966*; *Diez et al., 2007*), which are embedded in a dense exopolymeric matrix covering the surface and endolithic pore space of the beachrock (*Petrou et al., 2014*). We previously demonstrated the presence of Chl *f*-containing cyanobacteria (*Chroococcidiopsis* spp.) in an endolithic niche below the surface biofilm along with the ability of beachrock samples to upregulate their Chl *f*-content upon incubation under NIR, indicative of FaRLiP (*Trampe and Kühl, 2016*). Here, we correlate the spatial organization of Chl *f*-containing cyanobacteria with direct *in vivo* measurements of their NIR-driven O_2_ production in a natural beachrock habitat.

## Results and Discussion

Hyperspectral reflectance imaging on vertical cross-sections of beachrock submerged in seawater (23°C and salinity = 35) revealed the presence of a dense ~1 mm thick surface biofilm with high amounts of Chl *a*, while a more patchy zone containing Chl *f*, and less Chl *a* was found below the surface biofilm of the beachrock (Fig. 1A, B), exhibiting localized hot spots of Chl *f* concentration (Fig. 1C). Representative reflectance spectra from these regions carrying spectral signatures of maximal Chl *a* and Chl *f* absorption at 670–680 nm and 715–725 nm, respectively, are presented in the Supplementary Materials (Fig. S1) along with additional examples of hyperspectral imaging of beachrock cross-sections (Fig. S2). To further describe the microscale distribution of cells with different photo-pigmentation, we employed hyperspectral fluorescence imaging with confocal laser scanning microscopy (CLSM; 488 nm excitation) on beachrock cross-sections (Figs. S3, S4). The CLSM data confirmed the occurrence of patches of Chl *f*-containing cyanobacteria with a characteristic fluorescence peak around 740-750 nm (cf. *20*) in deeper endolithic zones (Figs. S3C, S4A-D). Brightfield microscopy of the Chl *f* hot spots, revealed the presence of round cell aggregates (Fig. S4E, F) typical of pleurocapsalean cyanobacteria (*Waterbury and Stanier, 1978*). These data confirm earlier findings of Chl *f* (and only minor amounts of Chl *d*) in beachrock associated to endolithic, *Chroococcidiopsis*-like cyanobacteria (*Trampe and Kühl, 2016*), and observations of Chl *f* in deeper, shaded layers of terrestrial epilithic biofilms (*Behrendt et al. 2015*) and microbial mats (*Ohkubo and Miyashita, 2017*).

**Figure 1.**
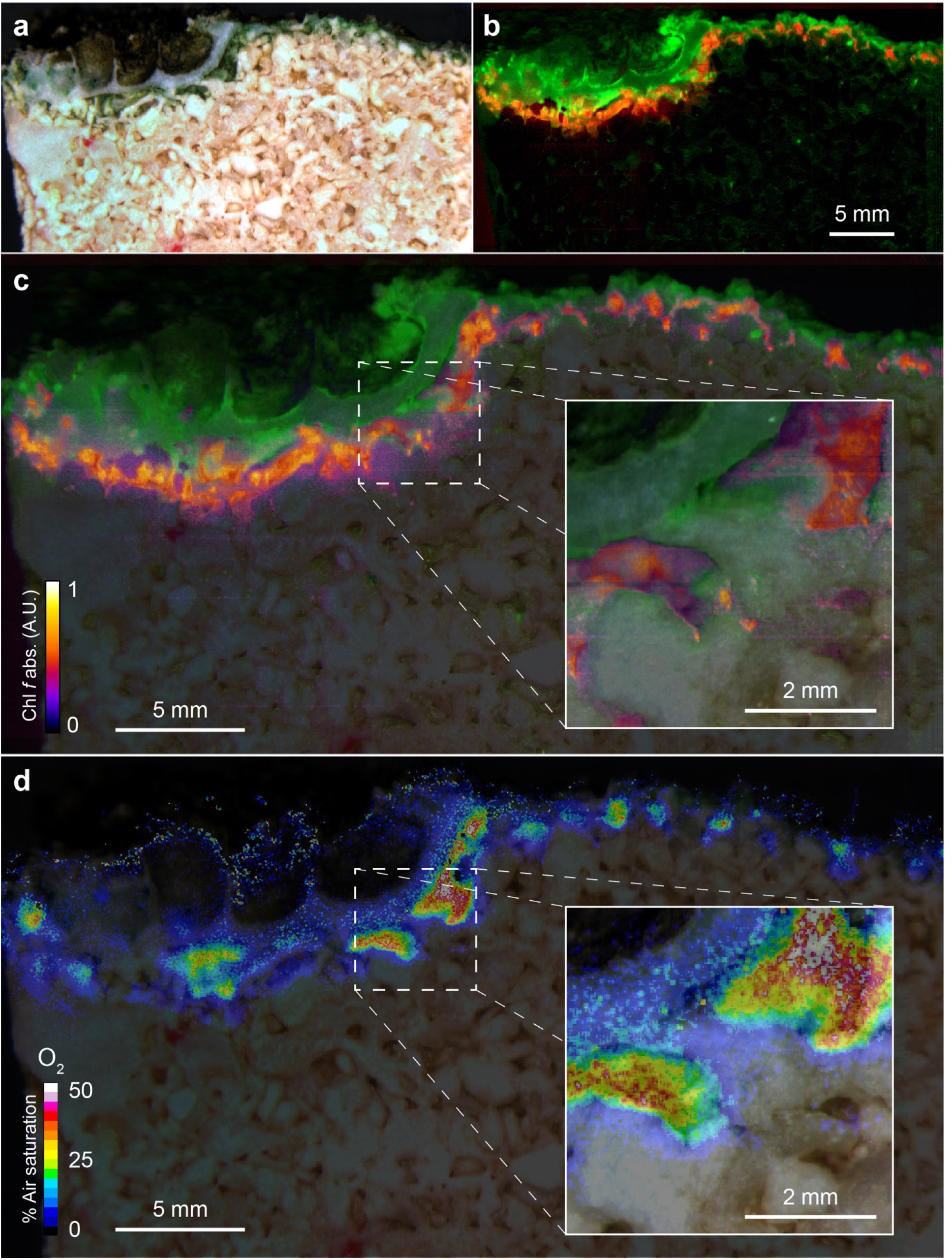
Spatial distribution of photopigments and near-infrared radiation driven oxygenic photosynthesis in beachrock as mapped with hyperspectral reflectance imaging and chemical imaging of O_2_. (A) RGB image composite, constructed from the hyperspectral image stack (R=650 nm, G=550 nm, B=450 nm), showing “true” colors of beachrock material and the biofilm community in a cross-section of the top layer. (B) False color coded image of the same hyperspectral image stack as in panel A mapping pixels with Chl *a* absorption (670-680nm) in green, and Chl *f* absorption (718-722nm) in red. Representative reflectance spectra of the two regions are given in Fig. S1. (C) Overlay of beachrock structure obtained in panel A and the Chl *a* signature from panel B with map of the relative abundance of Chl *f* obtained from the amplitude of Chl *f* absorption (color coded between 0 and 1), as acquired from hyperspectral image analysis. (C) Distribution of O_2_ concentration (color coded in units of % air saturation) in the beachrock under illumination of 740 nm light (HBW = 25 nm; photon irradiance = 28 µmol photons m^−2^ s^−1^) when immersed in anoxic seawater, as imaged with the beachrock section covered with a thin paint of agarose containing O_2_–sensitive nanoparticles. The O_2_ concentration image was superimposed onto the structural image of the beachrock cross section. The insert is a digital zoom corresponding to the insert in panel C. Additional data on 2 other beachrock sections are available in the Suppl. Materials (Figs. S2, S6).

By coating beachrock cross-sections with a thin (<1 mm) layer of an O_2_ sensitive nanoparticle-agarose paint (see Materials and Methods and Fig. S5) and subsequent immersion in anoxic water, it was possible to map the local O_2_ production over the beachrock cross-section when illuminated with weak NIR levels (740 nm, 25 nm HBW; 28 µmol photons m^−2^ s^−1^). We observed hot spots of NIR-driven photosynthesis driving local O_2_ levels from 0% to >40-50% air saturation within 15-20 minutes (Fig. 1D, SFig. S6), which overlapped with regions of high Chl *f* absorption (Fig. 1C, Fig. S2). The build-up of O_2_ in the hotspots harboring Chl *f* occurred rapidly after onset of NIR illumination and dissipated rapidly back to anoxia within a few minutes after darkening (see Movie S1). Based on O_2_ concentration images recorded at 5 min intervals after experimental light-dark shifts, we calculated images of apparent dark respiration and NIR-driven net and gross photosynthesis that could be mapped onto the beachrock structure (Fig. 2A-D) showing that hotspots of activity aligned with the presence of Chl *f* (see Fig. 1B, C). We extracted estimates of maximum O_2_ conversion rates in particular regions of interest (ROI) showing high rates of NIR-driven gross photosynthesis of ~5-15 µmol O_2_ L^−1^ min^−1^ in the beachrock under the given actinic irradiance of 28 µmol photons m^−2^ s^−1^; a similar range was found for 2 other beachrock cross-sections (data not shown). These volume-specific rates fall among the upper range of maximal gross photosynthesis rates in aquatic phototrophs, and are comparable with photosynthetic rates found in benthic microalgae (*Krause-Jensen and Sand-Jensen, 1998*).

**Figure 2.**
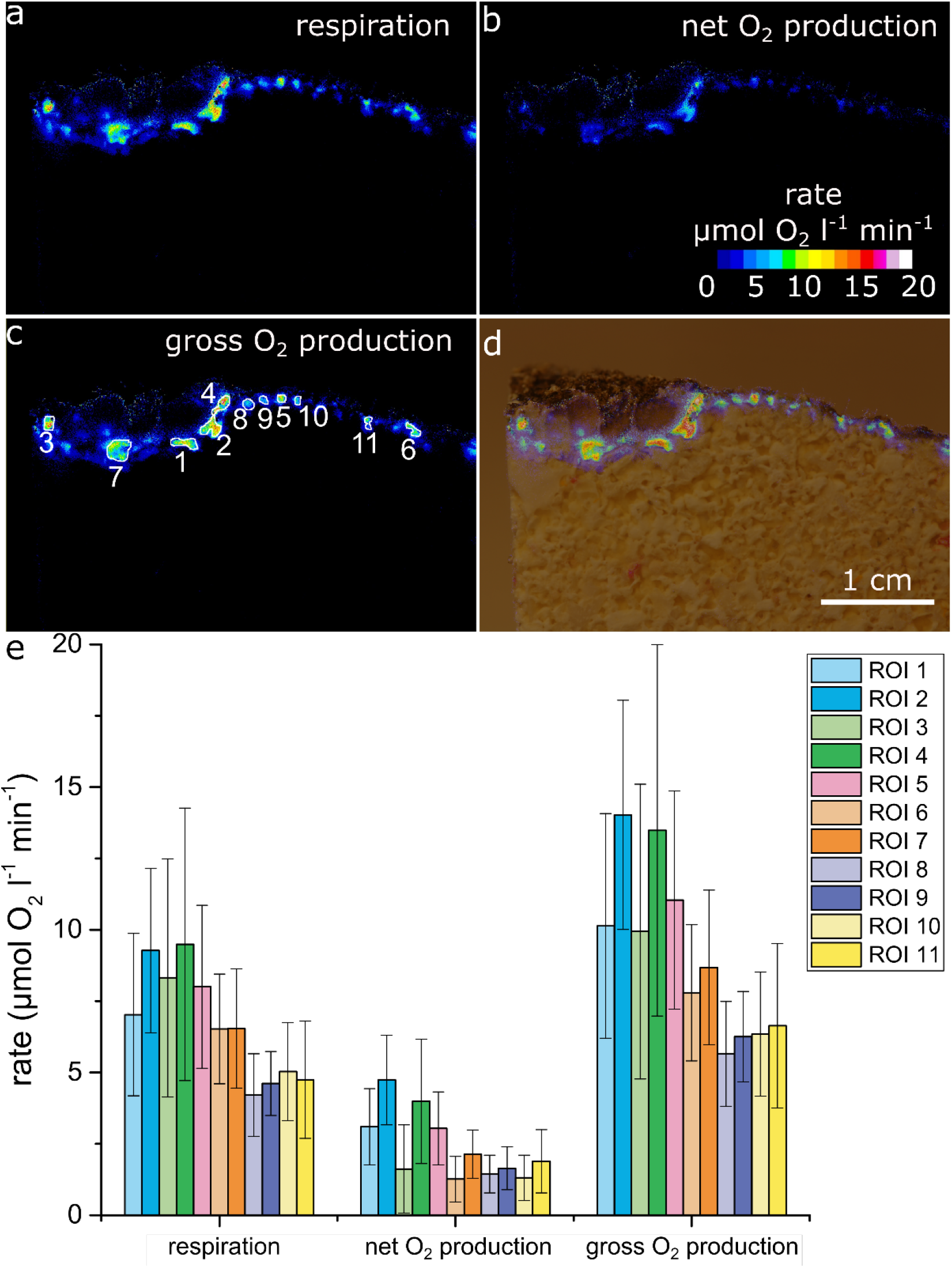
Oxygen consumption and NIR-driven oxygenic photosynthesis in beachrock. Cross-sectional images of initial O_2_ consumption after onset of darkness (A) and maximum net photosynthetic O_2_ production (B) after onset of actinic NIR illumination (740 nm, 28 µmol photons m^−2^ s^−1^) of the beachrock cross-section shown in Fig.1. (C) The NIR-driven gross photosynthesis was estimated by summing the absolute rates of net photosynthesis under NIR and O_2_ consumption in the dark. (D) Overlay of gross photosynthesis distribution over a structural image of the beachrock cross-section. (E) Data for O_2_ consumption, and NIR-driven net and gross photosynthesis were extracted for 11 regions of interest (ROI) in panel C and are presented as means± standard deviation within the ROI.

The Chl *f*-driven oxygenic photosynthesis by endolithic cyanobacteria thus seems very efficient given the low actinic NIR level applied. High photosynthetic efficiency of Chl *f*-containing cyanobacteria under NIR has been shown in previous *ex situ* studies on cultivated strains and enrichments. Gan et al. (2014) thus found that *Leptolyngbya* strain JSC-1 cells showed a 40% higher O_2_ production rate with NIR after undergoing FaRLiP relative to cells grown under red light (*Gan et al., 2014*), and a similar high photosynthetic efficiency was found in a *Chroococcidiopsis* strain (*Nürnberg et al., 2018*). Behrendt et al. (2015) showed rapid saturation of NIR-driven oxygenic photosynthesis already at 25-30 µmol photons m^−2^ s^−1^ (740 nm) in a cell enrichment with Chl *f*-containing, *Aphanocapsa*-like cyanobacteria from a cavernous biofilm, i.e., using similar NIR levels as used in the present study.

Our structural and chemical imaging of beachrocks showed that Chl *f* and NIR-driven O_2_ production was confined to a relative narrow zone 1-2 mm below the beachrock surface (Figs. 1, 2; Figs. S2–S5). Assuming a NIR-driven oxygenic photosynthesis rate of 10 µmol O_2_ L^−1^ min^−1^ (= nmol O_2_ cm^−3^ beachrock min^−1^) in a 1 mm thick layer (Fig. 2), and a conservative estimate of beachrock porosity of ~0.4 in the uppermost 1-2 mm (*21*), we can estimate the areal NIR-driven gross photosynthesis rate to (10 µmol l^−1^ min^−1^ × 0.4 × 0.1 cm × 10^−6^ × 10000 × 60 =) ~0.24 mmol O_2_ m^−2^ beachrock h^−1^. Total beachrock primary productivity remains to be quantified in detail, but it is well known that beachrock habitats sustain high grazing rates of epifauna (*McLean, 1974; McLean, 2011*) and herbivorous reef fish (*Stephenson and Searles, 1960*).

To our knowledge, beachrock primary production has only been reported by Krumbein, who studied a Red Sea beachrock habitat (*Krumbein, 1979*). Based on his data on O_2_ exchange under water covered conditions (cf. Fig. 13 in *Krumbein, 1979*), we estimated a total gross photosynthesis of ~10 mmol O_2_ m^−2^ beachrock h^−1^. While based on several crude assumptions and comparing different habitats, these rough calculations indicate that NIR-driven photosynthesis could account for at least 2-3% of total areal photosynthesis in beachrock habitats. Furthermore, we note that we base this estimate on O_2_ dynamics over minutes measured in a thin agar layer adjacent to the actual phototrophic cells, which will underestimate the true dynamics (see Methods), and our measurements were conducted at low NIR levels that may not represent saturated photosynthetis rates. Obviously, the realized total daily primary productivity will also be strongly modulated by the actual biomass distribution, porosity and diel light exposure within the beachrock, as well as the pronounced diel cycles of environmental conditions on the beachrock platform (*Petrou et al., 2014*; *Schreiber et al. 2002*). Nevertheless, we argue that our experimental data point to a significant role of NIR-driven oxygenic photosynthesis in beachrock and potentially other endolithic habitats.

Based on the origin of isolates shown to employ FaRLiP (Table S1), it has been speculated that shaded soils, caves, plant canopies and thermal springs may be prime terrestrial habitats for cyanobacteria with Chl *f* (*Gan et al. 2015*; *Zhang et al. 2019*). Beachrock is widespread in intertidal zones on a global scale (*Vousdoukas et al., 2007*) and our study shows that these habitats may present a major ecological niche for cyanobacteria with FaRLiP and NIR-driven oxygenic photosynthesis in marine intertidal habitats. The present study gives the hitherto most detailed insight into the distribution and *in situ* activity of Chl *f*-containing cyanobacteria in intact samples from a natural habitat, but there is now a need for more precise characterization of the microenvironment and metabolic activities of microorganisms in beachrock to assess the quantitative importance of NIR-driven oxygenic photosynthesis for system productivity.

Besides their strong relevance for exploring the biophysical and biochemical limits and controls of oxygenic photosynthesis, cyanobacteria with Chl *d* and Chl *f* have been employed in speculations on more efficient light harvesting of solar energy, as organism with red-shifted photopigments can in principle exploit ~20% more photons in the solar spectrum for their photosynthesis (*Chen and Blankenship, 2011*). However, FaRLiP involves remodeling of the photosynthetic apparatus for better performance under 700-760 nm light and is apparently not induced in absence of NIR (*Gan et al., 2014*), and *Acaryochloris marina* has largely exchanged its Chl *a* with Chl *d* (*Myashita et al., 2014*). Hence, the selective benefit of employing Chl *d*, Chl *f*, and red-shifted phycobilins in oxygenic photosynthesis appears more related to the capability of exploring special ecological niches in the shadow of other oxygenic phototrophs (*Ohkubo and Miyashita, 2017*; *Kühl et al., 2005*). The present study and the few other studies of natural habitats (Table S1) demonstrate that a key trait of such cyanobacteria is the formation of biofilms in strongly shaded environments below other algae, cyanobacteria or terrestrial plants, or in the twilight zone of caves (*Behrendt et al., 2019*). It remains to be explored to what extent the presence of cyanobacteria with FaRLiP-capability or constitutive high Chl *d* levels play a role for overall photosynthetic productivity in such habitats. That is, does the complementary photopigmentation of cyanobacteria with far-red shifted photopigments in the “understory” of other oxygenic phototrophs lead to higher photosynthetic efficiency and productivity at the community/system level, e.g. in analogy to plant canopies? Such studies are complicated due to the compacted and often stratified structure of the natural communities, but the novel combination of structural and chemical imaging presented here seems a promising toolset for unraveling the ecological importance of FaRLiP and cyanobacteria with far red-shifted photopigments.

## Materials and Methods

### Field site and beachrock sampling

Beachrock fragments (5-10 cm) from the upper black-brown zone of the intertidal beachrock platform on Heron Island (Great Barrier Reef, Queensland, Australia; 23°26.5540S, 151°54.7420E) were sampled at low tide and cut into smaller subsamples with smooth vertical cross sections (ca. 2 × 2 × 1 cm^3^) using a seawater-flushed stone saw under dim light. The samples were subsequently stored and transported dry and dark prior to experiments at University of Technology Sydney, which commenced within a few days after sampling. The field site is described in detail elsewhere (*Cribb, 1966*; *Diez et al., 2007*; *Trampe and Kühl, 2016*).

### Hyperspectral imaging

#### Imaging setup

Hyperspectral reflectance imaging was done on vertical cross-sections of beachrock as previously described (*Trampe and Kühl, 2016*). The samples were submerged in seawater (23°C and salinity = 35) with the cross-section facing the objective of a dissection zoom microscope (Leica, Germany) with a hyperspectral camera system (100T-VNIR; Themis Vision Systems, St. Louis, MO, USA) (*Kühl and Polerecky 2008*) connected via the C-mount video adapter. A fiber-optic halogen lamp with an annular ring-light (KL-2500 and Annular Ring-light; Schott AG, Mainz, Germany) mounted on the objective of the dissection microscope was used as a light source for the hyperspectral image acquisition. Additional hyperspectral measurements of reflected light were performed (at similar distance and microscope settings as used for the beachrock samples) on a calibrated 20% reflectance standard (Spectralon SRM-20; LabSphere Inc., North Sutton, NH, USA).

#### Hyperspectral image analysis

Using the manufacturers software, (PhiLumina Hyperspectral Imager V. 4.2; PhiLumina, LLC, Gulfport, MS, USA), dark-corrected, hyperspectral image stacks of beachrock reflection were converted to hyperspectral reflectance images (in units of % reflectance) by normalization to dark-corrected reflection stacks recorded using the reflectance standard. Subsequent file format conversion, and image cropping were performed in ENVI (Exelis Visual Information Solutions, Boulder, CO, USA), before further image processing with the software Look@MOSI (freeware available at www.microsen-wiki.net/doku.php/hsimaging:hs_iman_howto).

RGB images were constructed from the reflectance measurements at 650 nm (Red), 550 nm (Green), and 450 nm (Blue), as extracted from the calibrated hyperspectral image stacks. The fourth derivative of the hyperspectral reflectance stacks was calculated using the Look@MOSI software, yielding the relative extent of light attenuation at wavelengths representative for Chl *a* (670–680 nm) and Chl *f* (718–722 nm) absorption. The resulting greyscale images were then used for construction of false-color-coded images, showing the spatial coverage of the two defined spectral signatures. The extraction of spectral information from areas of interest (covering hotspots of Chl *f*) was performed as previously described (*Polerecky et al., 2009*). The resulting images were first cropped to be of computational sizes for the Look@MOSI software, and were stitched back together in Photoshop CC 2015.1.2 (Adobe Systems Incorporated, San Jose CA, USA) after computation. The relative absorbance/abundance of Chl *f* was quantified by scaling of the relative extent of light attenuation as calculated above for Chl *f.* Pixel intensity values from the earlier calculated grey scale images were assigned a scale ranging from 0 to 1, and were finally displayed using a color scaled lookup table in ImageJ (http://rsbweb.nih.gov/ij/). This yielded false-color coded images with values between 0 and 1, displaying a relative measure of Chl *f* abundance according to the amount of light attenuation obtained from Chl *f* absorption.

### Experimental setup for O_2_ imaging

The experiments were performed in a small glass aquarium (15 × 10 × 15 cm^3^) filled with filtered seawater (salinity 35). A lid with two inlets was placed on the aquarium. One inlet was used as a gas inlet, the second inlet was used to insert a robust fiber-optic O_2_ sensor (OXR430 connected to FireSting GO2 meter; PyroScience GmbH, Aachen, Germany) for monitoring the O_2_ concentration in the bulk seawater. The beachrock sample was mounted with a smooth vertical cross section parallel to the aquarium glass wall. The camera used for O_2_ imaging and the excitation LED were placed perpendicular to the sample cross section. Actinic NIR illumination was provided by a 740 nm LED (LZ4-40R300; LED Engin, Inc., San Jose, CA, USA; HBW 25 nm) providing a NIR photon irradiance (integrated over 715-765 nm) of 28 µmol photons m^−2^ s^−1^, as measured with a calibrated spectroradiometer (MSC15, GigaHertz-Optik GmbH, Germany).

A beachrock sample painted on one side with the O_2_ sensor nanoparticle paint (see below and Fig. S5) was placed into the aquarium, and the surrounding seawater was flushed with pure N_2_ for at least 30 min to completely remove O_2_, as confirmed by the fiber-optic O_2_ sensor. After anoxic conditions were reached, measurements of the change in O_2_ concentration over the beachrock cross section in darkness and under NIR illumination were performed under stagnant anoxic conditions in the surrounding seawater (Fig. 1C, Fig. S6). Images were recorded at 5 min intervals relative to onset of actinic light or darkness. This enabled both recording of steady-state O_2_ images in light and darkness (=homogeneous anoxia), as well as the dynamic change in O_2_ distribution upon NIR irradiation. Proxies for the NIR-driven net photosynthesis, P_N_, and apparent dark respiration, R_D_, were calculated by subtracting images taken with a 5 min interval just before and after onset or eclipse of the actinic NIR light, respectively. Gross photosynthesis was estimated as P_G_=P_N_+ǀR_D_ǀ. To avoid any interference from surrounding light reaching the sample, the entire setup was covered thoroughly with black fabric. All measurements were performed at room temperature (23°C ± 1°C).

It should be noted that the O_2_ measurements were done over minute intervals in the thin agar layer with O_2_ sensitive nanoparticles coating the beachrock with the photosynthetic cells, where the observed O_2_ levels are affected both by cell activity and diffusional exchange with the surroundings. This leads to some diffusive smearing, and the observed dynamics in O_2_ and thus the derived reaction rates likely represent an underestimate of “true” reaction rates (*Santner et al., 2015*).

### Chemical imaging of O_2_

#### Imaging system and application of O_2_ sensor nanoparticles

We used recently published protocols for imaging O_2_ concentration over complex bioactive surfaces by coating beachrock cross-sections with O_2_-sensitive luminescent sensor nanoparticles (*Koren et al., 2015*; *Koren et al. 2016*) followed by ratiometric luminescence imaging of the coated surface with a DSLR camera (without NIR filter and with a 530 nm long pass filter mounted on the camera objective) during brief excitation with blue LED light (445 nm) Details of the imaging system and image acquisition software (*Larsen et al., 2011*), nanoparticle fabrication (*Koren et al., 2015*; *Mosshammer et al., 2019*) and biocompatibility (*Trampe et al., 2018*) are given elsewhere.

#### Optical O_2_ sensor nanoparticle paint

First, we attempted to spray-paint beachrock sections with sensor nanoparticles using a paintbrush according to Koren et al. (2016), but the porous structure of the beachrock prevented saturation and a homogenous coating of cross-sections. Instead, we coated beachrock cross-sections (as well as glass slides used for calibration) with a thin (<1 mm thick) layer of agarose containing O_2_ sensor nanoparticles, inspired by earlier work on seagrass rhizosphere O_2_ imaging (*37*). For this, 40 mg of UltraPure™ Low Melting Point Agarose (16520100; thermofisher.com) was melted in 2 mL filtered seawater, which was then kept at ~35°C and mixed with 2 mL of a pre-warmed (~35°C) O_2_ sensor nanoparticle solution (2.5 mg mL^−1^). This “sensor paint” was then applied on a vertical cross-section of a beachrock sample (or a glass microscope slide) using a small paint brush. After solidification of the sensor paint, the coated beachrock was transferred into the aquarium and left there prior to experiments to allow acclimatization. Throughout sample preparation, exposure to high light levels was avoided. This procedure produced a stable homogenous coating of beachrock cross-sections (Fig. S5) with an O_2_ sensing layer that could be easily peeled off, enabling the same sample to be used for subsequent CLSM or hyperspectral imaging

#### Imaging and image analysis

The O_2_-dependend red emission, and the constant green reference emission from the sensor nanoparticles in the paint during brief excitation pulses from a blue LED (445 nm) were recorded in RGB pictures with a DSLR camera system (*Larsen et al,. 2011*) imaging coated beachrock sections or coated microscope glass slides. Mapping of NIR-driven oxygenic photosynthesis was done by O_2_-imaging of beachrock biofilm cross-sections immersed in anoxic (N_2_ flushed) seawater under illumination with a NIR LED. Acquired RGB images were split into red, green, and blue channels and analyzed using the freely available software ImageJ (http://rsbweb.nih.gov/ij/). First, the red channel images (recording the O_2_ sensitive emission of the sensor nanoparticles) and the green channel images (recording the constant emission of a reference dye in the nanoparticles) were divided using the ImageJ plugin Ratio Plus (http://rsb.info.nih.gov/ij/plugins/ratio-plus.html) in order to get ratio images, R (= red channel/ green channel).

Background fluorescence from Chl *a* (at >680 nm) excited by the excitation light (445 nm) during O_2_ imaging can potentially overlap with the red channel. Furthermore, such Chl *a*-based photosynthesis may also generate a small amount of O_2_ during the brief excitation pulse. To account for such potential artefacts, and as we aimed to visualize changes in O_2_ concentration (ΔO_2_) attributed to Chl *f*-based photosynthesis, all ratio images were subtracted from ratio images recorded after 45 min in the dark in anoxic water, R_dark_, (ΔR=R_dark_-R). Further, to avoid any cross-talk of NIR into the red channel, O_2_ images were always recorded during a brief (<1 s) darkening during image acquisition.

#### Calibration

For calibration, a glass slide was coated with a thin layer of the O_2_ sensor nanoparticle paint. TH coated slide was placed in the experimental setup with aerated seawater and imaged at identical camera settings as used for beachrock sample imaging. Subsequently, the O_2_ content of the seawater was decreased by flushing it intermittently with N_2_ gas under constant monitoring by a fiber-optic O_2_ microsensor (OXR430 connected to FireSting GO2 meter; PyroScience GmbH, Aachen, Germany). Calibration curves were obtained from RGB images of the calibration target recorded under a series of known seawater O_2_ concentrations (ranging from 100% air saturation to anoxia) using a ROI covering the whole field of view. Plotting R versus O_2_ showed an exponential decay with increasing O_2_ concentration (Fig. S7), as commonly observed for optical O_2_ sensing materials (*Koren and Kühl, 2018*; *Mosshamme et al., 2019*). To enable calibration of background corrected experimental ratio images (see above), we generated a calibration curve by plotting ΔR (=R_anoxic_-R) versus O_2_ concentration (Fig S7B), which was fitted with an exponential function in Image J. The ΔR images from the experiments were then calibrated in Image J using the exponential fit in the calibration function.

### Confocal laser scanning microscopy

After the O_2_-imaging, the nanoparticle paint was peeled off the beachrock sections, and a smaller subsection of the flat beachrock cross section (previously used for hyperspectral or chemical imaging) was imaged in a 35 mm coverslip glass bottom petri dish (World Precision Instruments) (with the beachrock cross section facing downwards) at 200x and 400x magnification on an inverted confocal laser scanning microscope (Nikon A1R, Japan). Care was taken to keep the beachrock cross sections intact and oriented to enable identification and alignment of CLSM data to areas of interest exhibiting NIR-driven O_2_ production.

The microscope was equipped with a motorized xyz sample holder and was able to acquire hyperspectral fluorescence image stacks (10 nm) over a large sample area by sequential scanning, with subsequent automatic stitching of the acquired images in the microscope software (NIS elements AR, Nikon, Japan). The sample was excited by 488 nm laser light (laser power 1.2 mW, 0.5 frames per seconds) and spectral fluorescence (500-750 nm) was acquired at 10 nm resolution by the spectral PMT-array detector on the CLSM microscope. First, the beachrock cross-section was imaged at 200x magnification obtaining hyperspectral image stacks of 5 confocal layers (step size 5 µm) for each field of view. Scanned images were stitched to make a 4 mm × 11 mm large image and were then de-convolved (numerical aperture of 0.75, pinhole size 177.52, refractive index 1 for 26 emission channels 500-750 nm) using the NIS elements AR software (Version 4.60, Nikon, Japan). Subsequently, a more detailed scan of the same cross sectional area was done at 400x magnification (laser power 1.2 mW, 0.063 frames per seconds), and spectral fluorescence (500-750 nm) was acquired at 10 nm resolution by the spectral PMT-array detector on the CLSM microscope obtaining hyperspectral fluorescence images in one z-plane.

We note that the beachrock cross-section was not perfectly smooth and such single plane recording at 400x led to less signal in parts of the cross-section, where the surface was out of focus. Nevertheless, we could still identify most of the hot spots recorded at 200x magnification and resolve the shape of individual cells and cell clusters. Scanned images were stitched to make a 3.5 mm × 5.2 mm large image and were then de-convolved (numerical aperture of 1.00, pinhole size 177.52, refractive index 1.515 for 26 emission channels 500-750 nm). Regions of interest (ROI) were selected from the obtained cross-sectional scans and spectral fluorescence features were captured and false-color coded for particular regions of interest. CLSM scans were false-color coded to highlight the fluorescence of Chl *a* (690-700 nm), phycobiliproteins (650-660 nm), and Chl *f* (740-750 nm; cf. *20*).

## Acknowledgments

We acknowledge the excellent technical assistance in the field and during laboratory measurements by Sofie L. Jakobsen and the staff at Heron Island Research Station. Work at Heron Island was conducted under permit no. G16/38423.1 from the Great Barrier Reef Marine Parks authority

## Funding

This study was supported by project grants from the Independent Research Fund Denmark | Natural Sciences (DFF-8021-00308B; MK) & Technical and Production Sciences (DFF-8022-00301B and DFF-4184-00515B; MK), the Villum Foundation (MK) and the Poul Due Jensen Foundation (KK)

## Author contributions

MK, ET, KK designed the research, ET, KK, MM, AWD, MJ, MK conducted experiments, MJ, MK provided research infrastructure, ET, KK, MJ, MK analyzed data, and MK wrote the article with editorial input from all coauthors.

## Competing interests

The authors declare no competing interest.

## Data and materials availability

All data is available in the main text or the supplementary materials, and can be obtained from the corresponding author upon request.

## Supplementary Materials

**Figure S1.**
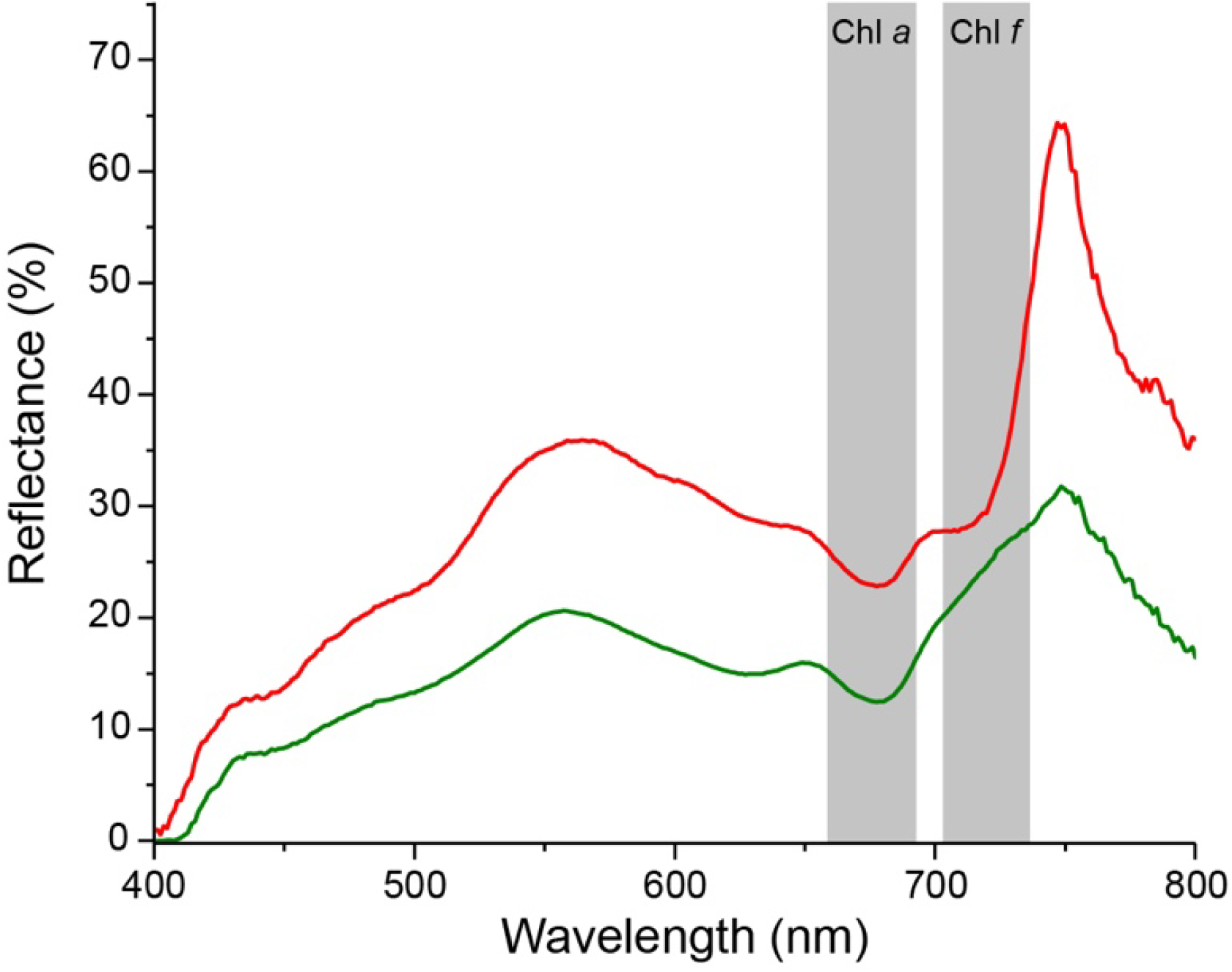
Representative reflectance spectra of beachrock zones with Chl *a* only (green zones in Fig. 1B, C) and zones with Chl *a* and Chl *f* (orange zones in Fig. 1B, C). The range of Chl *a* and Chl *f* absorption are indicated with shaded areas..

**Figure S2.**
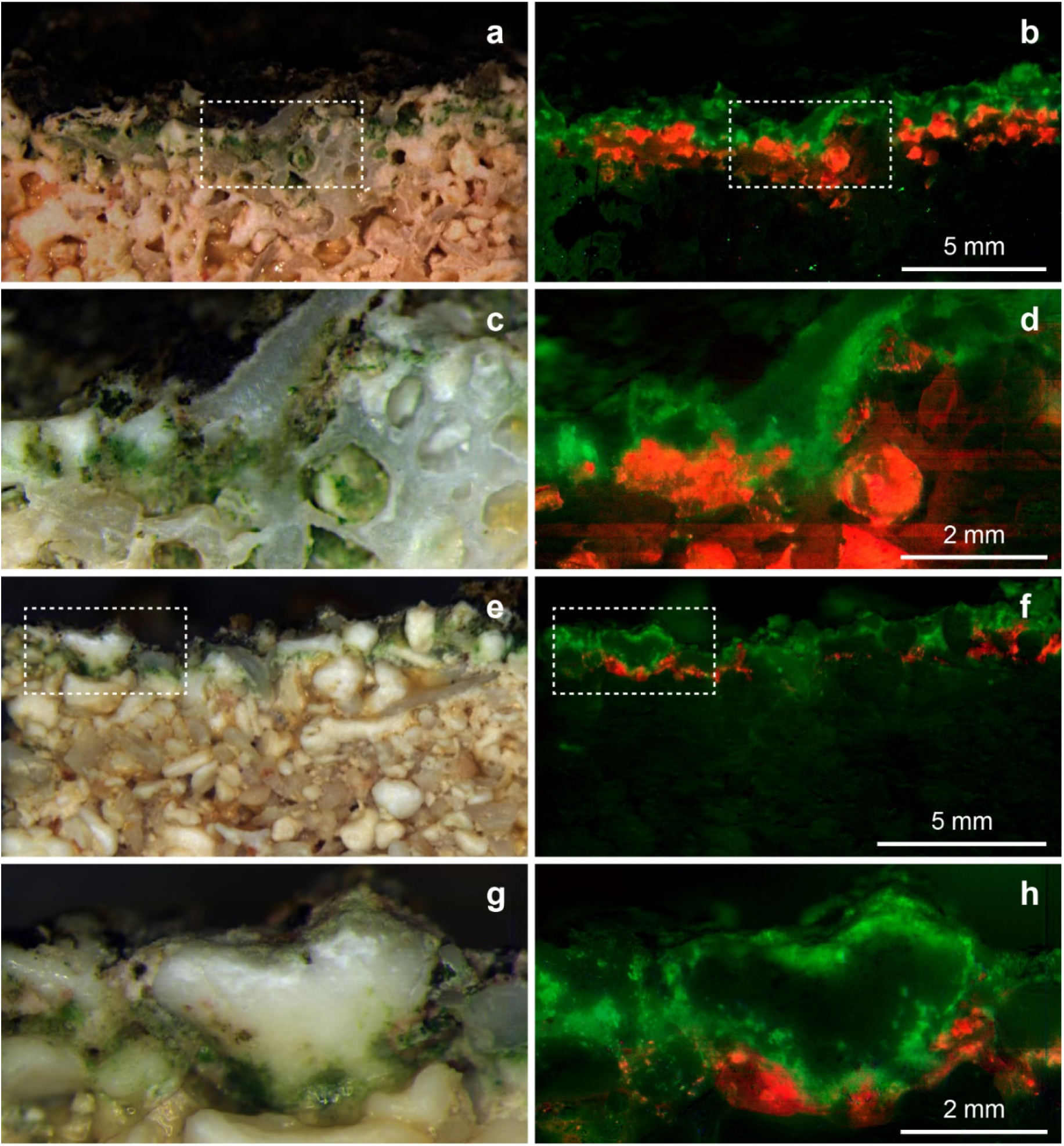
Macroscopic distribution of Chl *f* in beachrock Spatial distribution of photopigments as mapped with hyperspectral reflectance imaging in beachrock from the black colored (A-D) and brow-colored (E-F) zone on the beachrock platform on Heron Island. Left side panels; A, C, E, G are RGB composite images, constructed from the hyperspectral image stacks (R=650 nm, G=550 nm, B=450 nm), showing “true” colors of beachrock material and the biofilm community in a cross-section of the top layer. The right side panels; B, D, F, H are false color coding images of the corresponding hyperspectral image stacks adjacent to the left, they map pixels with Chl *a* absorption (675-680nm) in green, and Chl *f* absorption (718-722nm) in red. Representative reflectance spectra of the two regions are shown in Fig. S1C). Dashed boxes illustrate magnified parts displayed in the corresponding image directly below.

**Figure S3.**
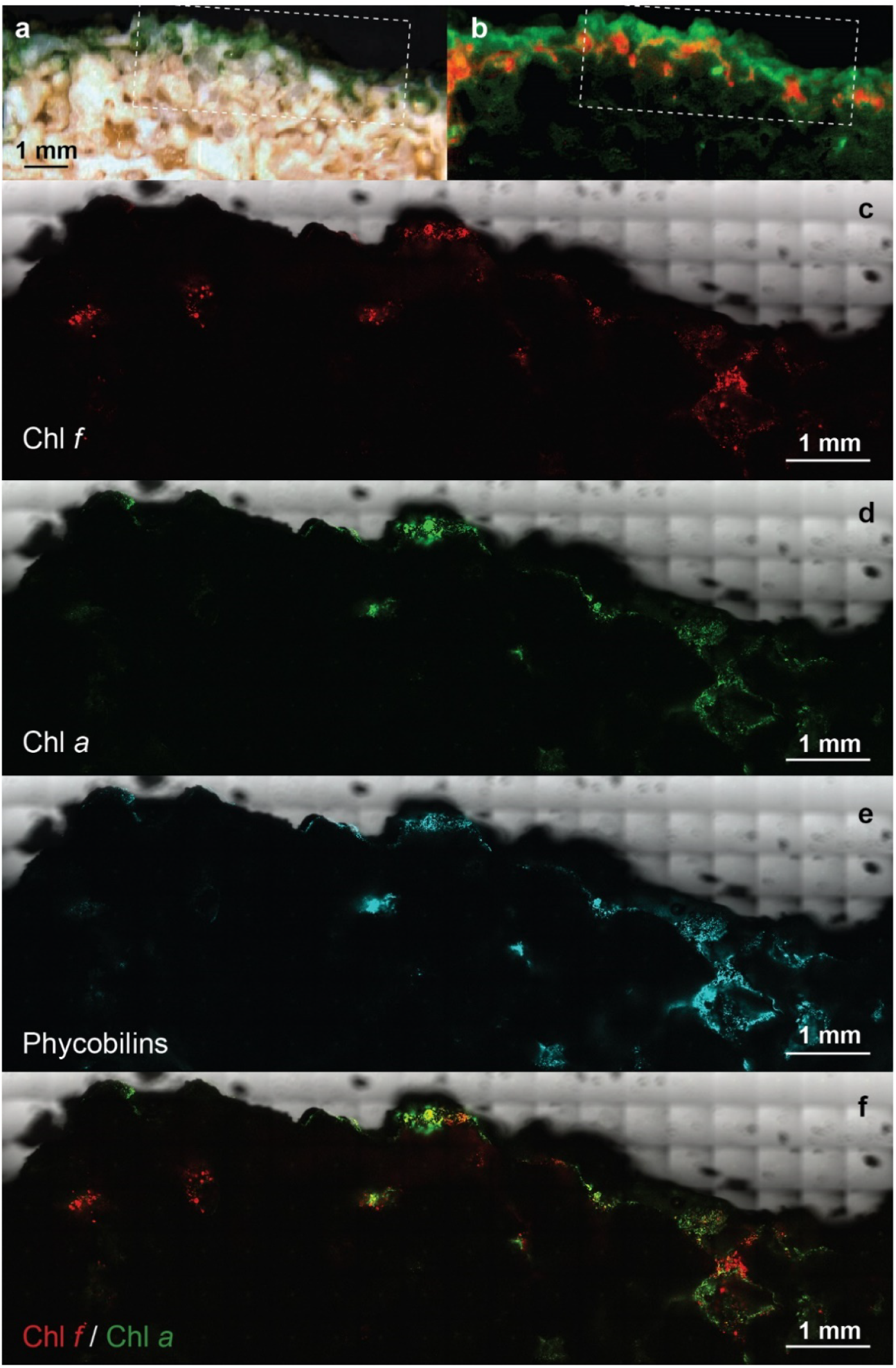
Confocal laser scanning microscopy (CLSM) of a beachrock cross-sectional area. (A) Hyperspectral RGB image, and (B) color-coded image showing Chl *a* (green) and Chl *f* (red) distribution over the same beachrock cross-section as in Fig. 1. Dotted boxes in panel A and B outline the area scanned with CLSM. Maximum projection image of pigment fluorescence over beachrock cross-sections imaged at 200x magnification for 5 focal planes (at 5 µm interval) highlighting areas with spectral fluorescence signatures of Chl *a* (C; 690-700 nm), phycobilins (D: 650-660 nm), Chl *f* (E; 740-750 nm), and the Chl *a* / Chl *f* ratio (F).

**Figure S4.**
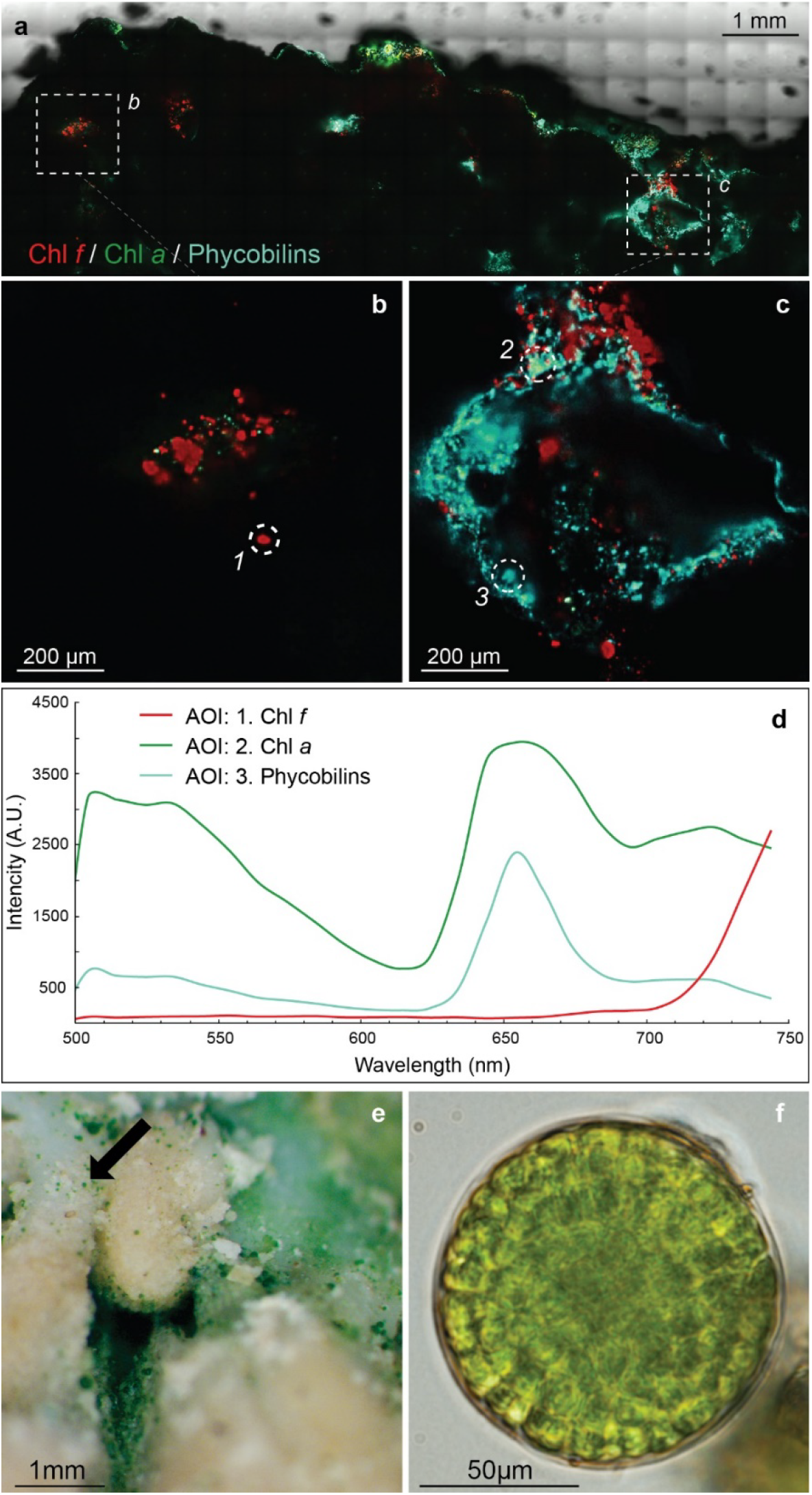
(A-C) Higher resolution CLSM scans of beachrock cross-section (marked with a dotted box in Fig S3A, B) recorded at 600x magnification. (D) False color coding of CLSM images were done based on spectral fluorescence characteristics of cells with Chl *a* (690-700 nm, green), phycobilins (650-660 nm, magenta), and Chl *f* (740-750 nm, red) in regions of interest (ROI) highlighted by circles in panel B and C. (E, F) Brightfield microscopy image of patches of green cell aggregates attached to beachrock (E) and a single cell aggregate of pleurocapsalean *Chroococcidiopsis*-like cyanobacteria retrieved from a ROI exhibiting characteristic Chl *f* fluorescence and absorption features (F).

**Figure S5.**
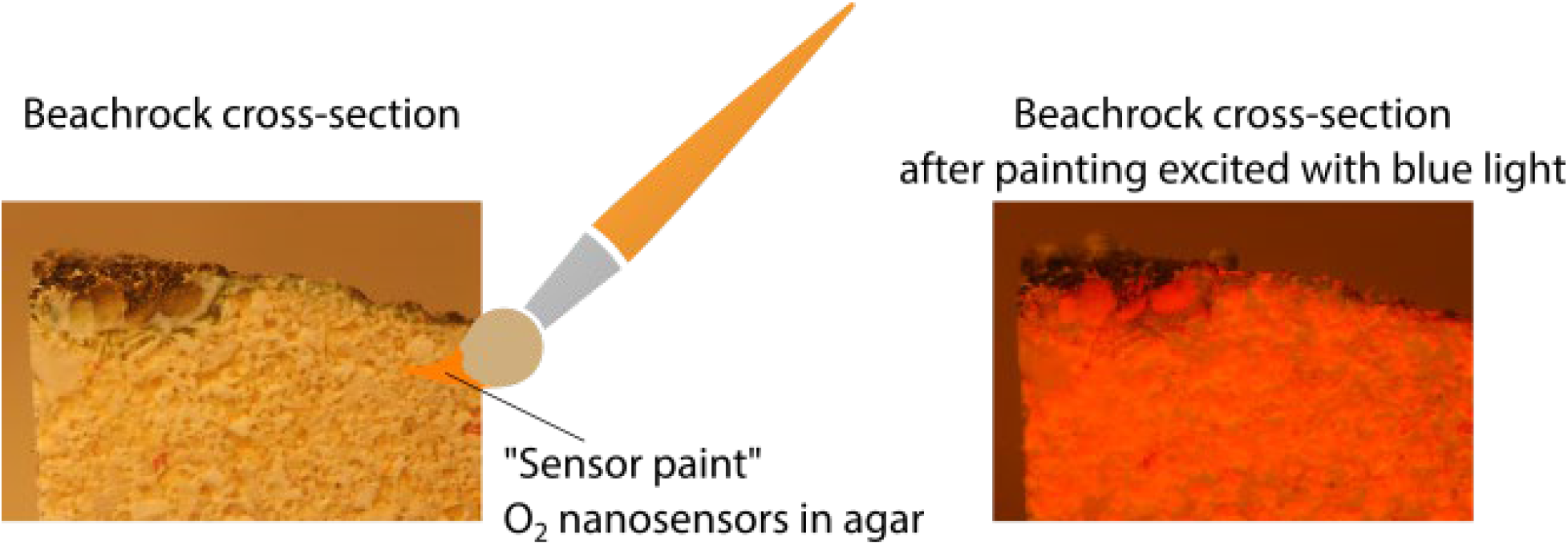
Application of agarose paint with nanoparticles Vertical cross-sections of beachrock (left panel) were painted with an agarose paint containing O_2_-sensitive luminescent nanoparticles. The beachrock was coated homogenously with a thin (< 1 mm) layer of luminescent paint (right panel) that was mechanically stable after solidification of the agar. This enabled ratiometric imaging of the O_2_ dependent luminescence over the beachrock cross-section (see Materials and Methods)

**Figure S6.**
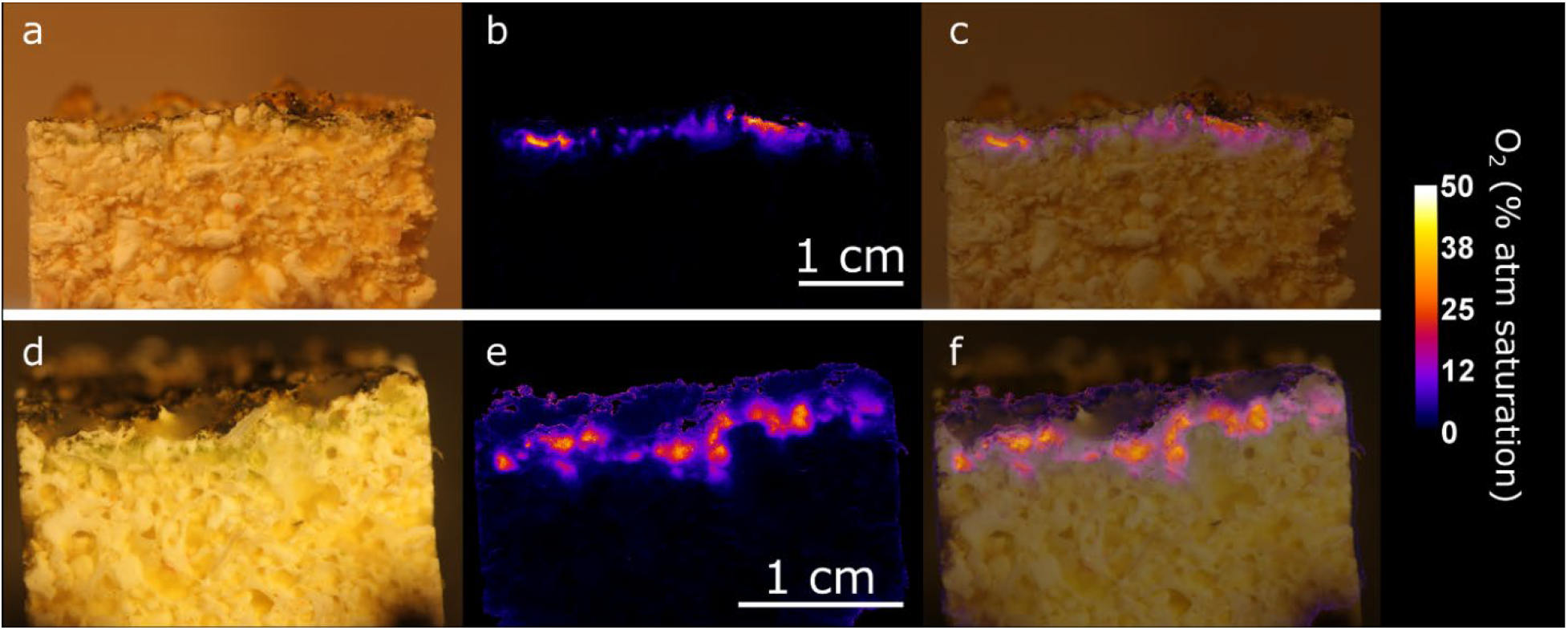
Additional examples of NIR-driven O_2_ production by endolithic cyanobacteria in beachrock. Cross-sections of 2 beachrock samples (A, D) were painted with an agarose paint containing O_2_ sensitive nanoparticles (see Fig. S5) and incubated in anoxic seawater for 30 minutes under a photon irradiance (740 nm, HBW 25 nm) of 28 µmol photons m^−2^ s^−1^. The resulting O_2_ concentration (false-color coded in units of % air saturation) was mapped via ratiometric luminescence imaging (B, E) and overlaid on top of the structural image of the beachrock cross-section (C, F).

**Figure S7.**
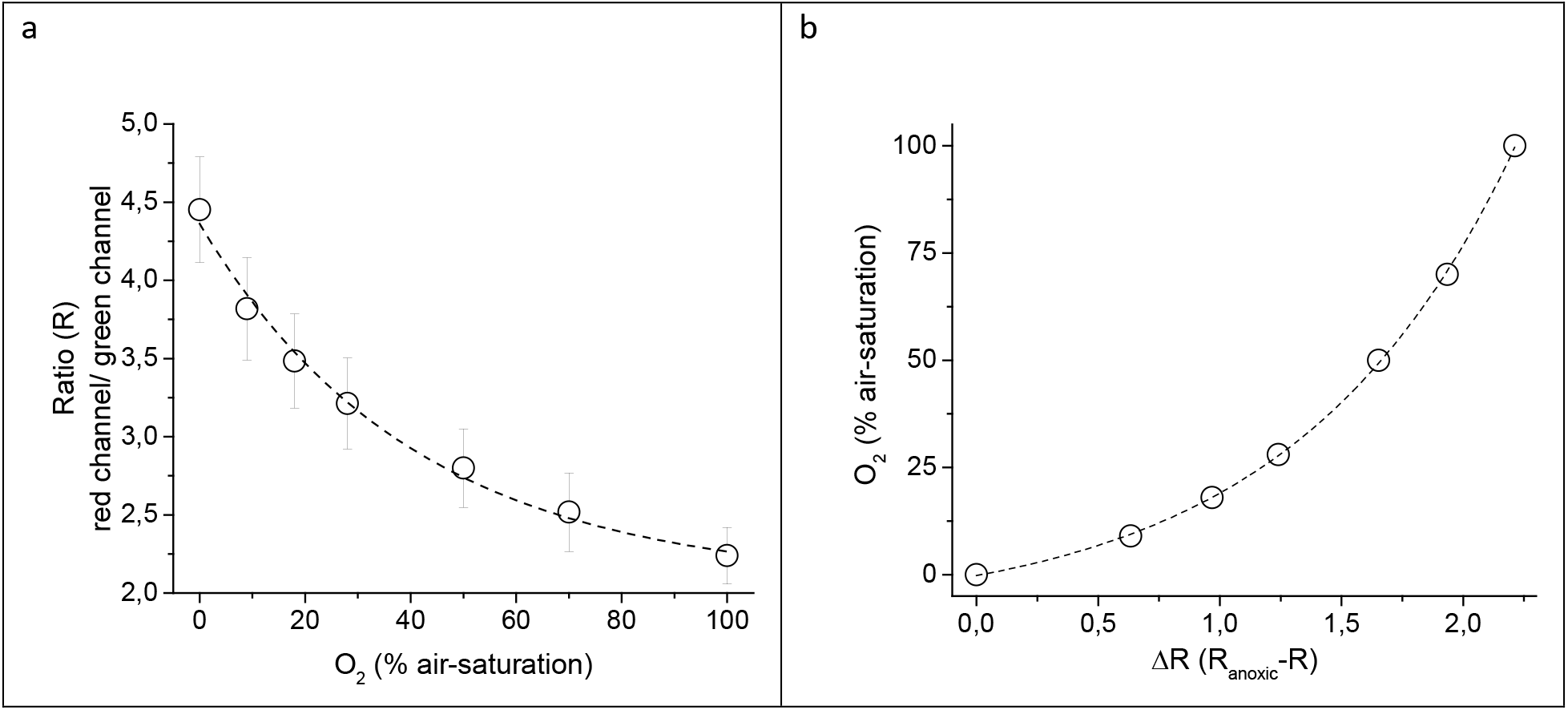
Calibration curve of the nanoparticle-based O_2_ sensor paint. (A) Ratio (R) of the red channel and the green channel in RGB images plotted vs. the measured O_2_ concentration in % air-saturation. (B) Calibration curve of the sensor particle based sensor paint calculated as ΔR (=R_anoxic_-R) plotted vs. the measured O_2_ concentration in % air-saturation. Data points with error bars represent means of the entire sensing area with the corresponding standard deviation; the dotted lines represent exponential fits (R^2^>0.99)

**Table S1.**
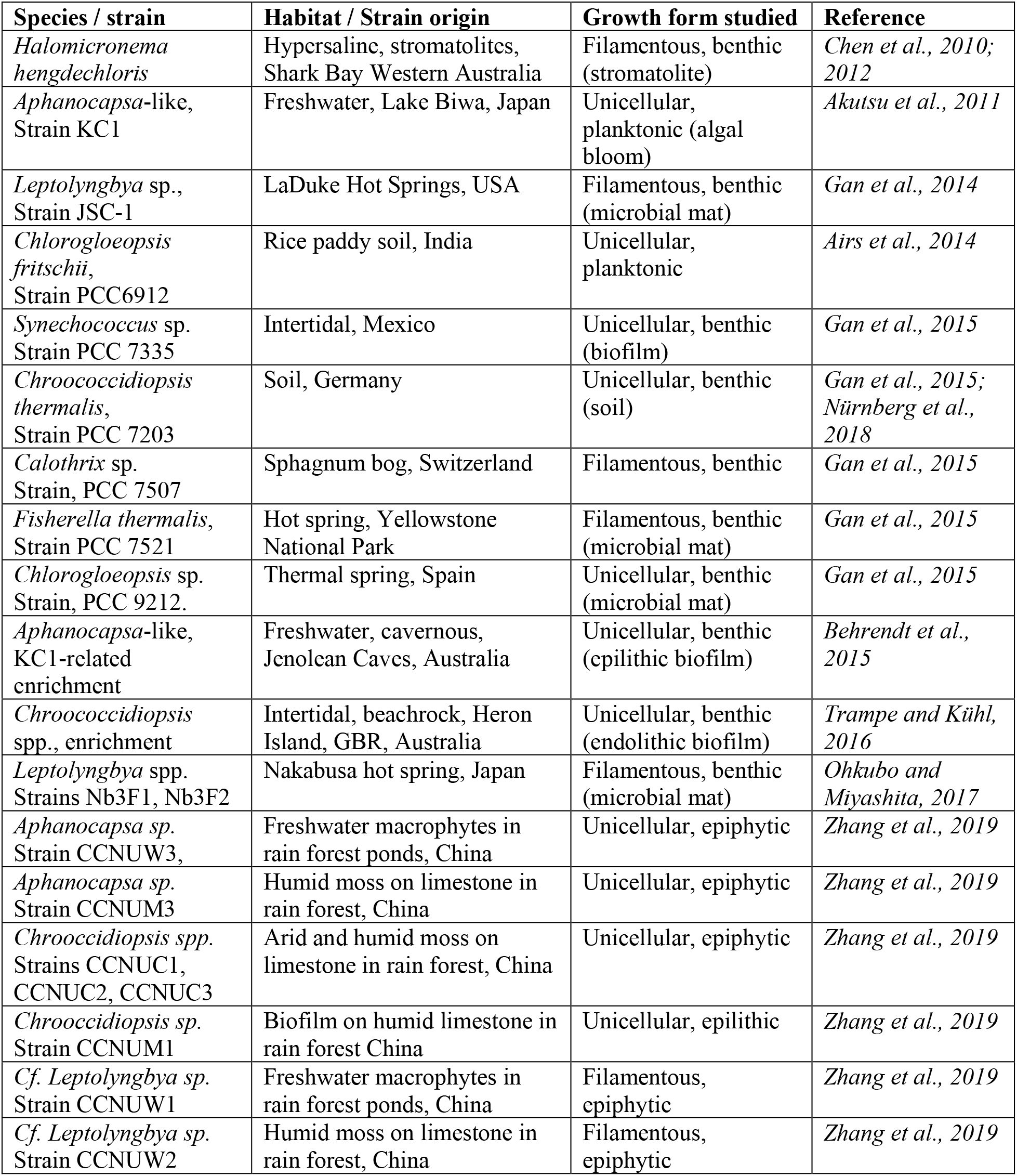
Overview of cyanobacterial strains and enrichments reported to contain Chl *f*. Listed in chronological order of first publication year.

| Species / strain | Habitat / Strain origin | Growth form studied | Reference |
| --- | --- | --- | --- |
| *Halomicronema hengdechloris* | Hypersaline, stromatolites, Shark Bay Western Australia | Filamentous, benthic (stromatolite) | *Chen et al., 2010; 2012* |
| *Aphanocapsa*-like, Strain KC1 | Freshwater, Lake Biwa, Japan | Unicellular, planktonic (algal bloom) | *Akutsu et al., 2011* |
| *Leptolyngbya* sp., Strain JSC-1 | LaDuke Hot Springs, USA | Filamentous, benthic (microbial mat) | *Gan et al., 2014* |
| *Chlorogloeopsis fritschii*, Strain PCC6912 | Rice paddy soil, India | Unicellular, planktonic | *Airs et al., 2014* |
| *Synechococcus* sp. Strain PCC 7335 | Intertidal, Mexico | Unicellular, benthic (biofilm) | *Gan et al., 2015* |
| *Chroococcidiopsis thermalis*, Strain PCC 7203 | Soil, Germany | Unicellular, benthic (soil) | *Gan et al., 2015; Nürnberg et al., 2018* |
| *Calothrix* sp. Strain, PCC 7507 | Sphagnum bog, Switzerland | Filamentous, benthic | *Gan et al., 2015* |
| *Fisherella thermalis*, Strain PCC 7521 | Hot spring, Yellowstone National Park | Filamentous, benthic (microbial mat) | *Gan et al., 2015* |
| *Chlorogloeopsis* sp. Strain, PCC 9212. | Thermal spring, Spain | Unicellular, benthic (microbial mat) | *Gan et al., 2015* |
| *Aphanocapsa*-like, KC1-related enrichment | Freshwater, cavernous, Jenolean Caves, Australia | Unicellular, benthic (epilithic biofilm) | *Behrendt et al., 2015* |
| *Chroococcidiopsis* spp., enrichment | Intertidal, beachrock, Heron Island, GBR, Australia | Unicellular, benthic (endolithic biofilm) | *Trampe and Kühl, 2016* |
| *Leptolyngbya* spp. Strains Nb3F1, Nb3F2 | Nakabusa hot spring, Japan | Filamentous, benthic (microbial mat) | *Ohkubo and Miyashita, 2017* |
| *Aphanocapsa* sp. Strain CCNUW3, | Freshwater macrophytes in rain forest ponds, China | Unicellular, epiphytic | *Zhang et al., 2019* |
| *Aphanocapsa* sp. Strain CCNUM3 | Humid moss on limestone in rain forest, China | Unicellular, epiphytic | *Zhang et al., 2019* |
| *Chroococcidiopsis* spp. Strains CCNUC1, CCNUC2, CCNUC3 | Arid and humid moss on limestone in rain forest, China | Unicellular, epiphytic | *Zhang et al., 2019* |
| *Chroococcidiopsis* sp. Strain CCNUM1 | Biofilm on humid limestone in rain forest China | Unicellular, epilithic | *Zhang et al., 2019* |
| *Cf. Leptolyngbya* sp. Strain CCNUW1 | Freshwater macrophytes in rain forest ponds, China | Filamentous, epiphytic | *Zhang et al., 2019* |
| *Cf. Leptolyngbya* sp. Strain CCNUW2 | Humid moss on limestone in rain forest, China | Filamentous, epiphytic | *Zhang et al., 2019* |

### Caption for Supplementary Movie S1

Animation of NIR-driven O_2_ dynamics over a beachrock cross-section (see Figs. 1 and 2) coated with a thin (<1 mm) agarose layer with luminescent O_2_ sensor nanoparticles. The movie sequence shows the decline in O_2_ concentration (recorded at 5 min interval) starting from steady-state conditions under a NIR irradiance (740 nm; 25 nm HBW) of 28 µmol photons m^−2^ s^−1^ approaching steady-state dark conditions after 35 min, followed by the rise in O_2_ concentration over 25 min after switching the NIR irradiation on again. The colored scale bar relates the colors to O_2_ concentrations.

## References

Airs RL, Temperton B, Sambles C, Farnham G, Skill SC, Llewellyn CA. 2014. Chlorophyll *f* and chlorophyll *d* are produced in the cyanobacterium *Chlorogloeopsis fritschii* when cultured under natural light and near-infrared radiation. FEBS Letters 588:3770–3777.

Akutsu S, Fujinuma D, Furukawa H, Watanabe T, Ohnishi-Kameyama M, et al. 2011. Pigment analysis of a chlorophyll *f*-containing cyanobacterium strain KC1 isolated from Lake Biwa. Photomedicine and Photobiology 33:35–40.

Allakhverdiev SI, Kreslavski VD, Zharmukhamedov SK, Voloshin RA, Korol’kova DV, Tomo T, Shen J-R. 2016. Chlorophylls *d* and *f* and their role in primary photosynthetic processes of cyanobacteria. Biochemistry 81:201–212.

Behrendt L, Trampe E, Larkum AWD, Qvortrup K, Norman A, Chen M, Ralph PJ, Sørensen SJ, Kühl M. 2011. Endolithic chlorophyll *d* containing cyanobacteria. ISME Journal 5:1072–1076.

Behrendt L, Brejnrod A, Schliep M, Sørensen SJ, Larkum AWD, Kühl M. 2015. Chlorophyll *f*-driven photosynthesis in a cavernous cyanobacterium. ISME Journal 9:2108–2111.

Behrendt L, Trampe EL, Nord NB, Nguyen J, Lonco D, Nyarko A, Dhinojwala A, Hershey OH, Kühl M, Barton H. 2019. Life in the dark: Far-red absorbing cyanobacteria extend photic zones deep into terrestrial caves. Environmental Microbiology DOI: 10.1111/1462-2920.14774.

Chen M. 2014. Chlorophyll modifications and their spectral extension in oxygenic photosynthesis. Annual Reviews in Biochemistry 83:317–340.

Chen M, Blankenship RE. 2011. Expanding the solar spectrum used by photosynthesis. Trends in Plant Science 16:427–431.

Chen M, Li Y, Birch D, Willows RD. 2012. A cyanobacterium that contains chlorophyll *f* — a red-absorbing photopigment. FEBS Letters 586:3249–3254

Chen M, Schliep M, Willows R, Cai Z-L, Neilan B, Scheer H. 2010 A red-shifted chlorophyll. Science 329:1318–1319.

Cribb AB. 1966. The algae of Heron Island, Great Barrier Reef, Australia, part I. A general account. University of Queensland Papers, Great Barrier Reef Commission, Heron Island Research Station 1:3–23.

Diez B, Bauer K, Bergman B. 2007. Epilithic cyanobacterial communities of a marine tropical beach rock (Heron Island, Great Barrier Reef): diversity and diazotrophy. Applied and Environmental Microbiology 73:3656–3668.

Gan F, Bryant DA. 2015. Adaptive and acclimative responses of cyanobacteria to far-red light. Environmental Microbiology 17:3450–3465.

Gan F, Shen G, Bryant DA. 2015. Occurrence of far-red light photoacclimation (FaRLiP) in diverse cyanobacteria. Life (Basel) 5:4–24.

Gan F, Zhang S, Rockwell NC, Martin SS, Lagarias C, Bryant DA. 2014. Extensive remodeling of a cyanobacterial photosynthetic apparatus in far-red light. Science 345:1312–1317.

Ho M-Y, Shen G, Canniffe DP, Zhao C, Bryant DA. 2016. Light-dependent chlorophyll *f* synthase is a highly divergent paralog of PsbA of photosystem II. Science 353:aaf9178.

Itoh S, Ohno T, Noji T, Yamakawa H, Komatsu H, Wada K, et al. 2015. Harvesting far-red light by chlorophyll *f* in photosystems I and II of unicellular cyanobacterium strain KC1. Plant & Cell Physiology 56:2024–2034.

Koren K, Kühl M. 2018. “Optical O_2_ sensing in aquatic systems and organisms” in Quenched-phosphorescence detection of molecular oxygen: Applications in life sciences, Papkovsky DB, Dmitriev R (Eds), Royal Society of Chemistry, Detection Science Series Volume 11, pp. 145–174.

Koren K, Jakobsen SL, Kühl M. 2016. In-vivo imaging of O_2_ dynamics on coral surfaces spray-painted with sensor nanoparticles. Sensors and Actuators B 237:1095–1101.

Koren K, Brodersen KE, Jakobsen SL, Kühl M. 2015. Optical sensor nanoparticles in artificial sediments – a new tool to visualize O_2_ dynamics around the rhizome and roots of seagrasses. Environmental Science & Technology 49:2286–2292.

Krause-Jensen D, Sand-Jensen K. 1998. Light attenuation and photosynthesis in aquatic plant communities. Limnology and Oceanography 43:396–407.

Krumbein WE. 1979. Photolithotropic and chemoorganotrophic activity of bacteria and algae as related to beachrock formation and degradation (Gulf of Aqaba, Sinai). Geomicrobiology Journal 1:156–202.

Kühl M, Polerecky L. 2008. Functional and structural imaging of phototrophic microbial communities and symbioses. Aquatic Microbial Ecology 53:99–118.

Kühl M, Chen M, Ralph PJ, Schreiber U, Larkum AWD. 2005. A niche for cyanobacteria containing chlorophyll *d*. Nature 433:820.

Larsen M, Borisov SM, Grunwald B, Klimant I, Glud RN. 2011. A simple and inexpensive high resolution color ratiometric planar optode imaging approach: application to oxygen and pH sensing. Limnology and Oceanography Methods 9:348–360.

Majumder ELW, Wolf BM, Liu H, Berg RH, Timlin JA, Chen M, Blankenship RE. 2017. Subcellular pigment distribution is altered under far-red light acclimation in cyanobacteria that contain chlorophyll *f*. Photosynthesis Research 134:183–192.

McLean RF. 1974. Geologic significance of bioerosion of beachrock. Proceedings of the Second International Coral Reef Symposium 2:401–408.

McLean RF. 2011. “Beach Rock” in Encyclopedia of Modern Coral Reefs, D. Hopley Ed., Encyclopedia of Earth Sciences Series (Springer, Dordrecht).

Miyashita H, Ohkubo S, Hirohisa K, Yuhta S, Daisuke F, Daiki F, et al. 2014. Discovery of chlorophyll *d* in *Acaryochloris marina* and chlorophyll *f* in a unicellular cyanobacterium, strain KC1, isolated from Lake Biwa. Journal of Physical Chemistry 4:1–9.

Mosshammer M, Brodersen KE, Kühl M, Koren K. 2019. Nanoparticle-based luminescence imaging of chemical species and temperature in aquatic systems: a review. Microchimica Acta 186:126.

Nürnberg DJ, Morton J, Santabarbara S, Telfer A, Joliot P, Antonaru LA, et al. 2018. Photochemistry beyond the red limit in chlorophyll *f*–containing photosystems. Science 360:1210–1213.

Ohkubo S, Miyashita H. 2017. A niche for cyanobacteria producing chlorophyll *f* within a microbial mat. ISME Journal 11:2368–2378.

Petrou K, Trimborn S, Kühl M, Ralph PJ. 2014. Desiccation stress in two intertidal beachrock biofilms. Marine Biology 161:1765–1773.

Polerecky L, Bissett A, Al-Najjar M, Faerber P, Osmers H, Suci PA, et al. 2009. Modular spectral imaging system for discrimination of pigments in cells and microbial communities. Applied and Environmental Microbiology 75:758–771.

Santner J, Larsen M, Kreuzeder A, Glud RN. 2015. Two decades of chemical imaging of solutes in sediments and soils – a review. Analytica Chimica Acta 878:9–42.

Schreiber U, Gademann R, Bird P, Ralph PJ, Larkum AWD, Kühl M. 2002. Apparent light requirement for activation of photosynthesis upon rehydration of desiccated beachrock microbial mats. Journal of Phycology 38:125–134.

Stephenson W, Searles RB. 1960. Experimental studies on the ecology of intertidal environments at Heron Island. I. Exclusion of fish from beach rock. Australian Journal of Marine and Freshwater Research 11:241–268.

Trampe E, Kühl M. 2016. Chlorophyll *f* distribution and dynamics in cyanobacterial beachrock biofilms. Journal of Phycology 52:900–996.

Trampe E, Koren K, Akkineni AR, Senwitz C, Krujatz F, Lode A, Gelinsky M, Kühl M. 2018. Functionalized bioink with optical sensor nanoparticles for O_2_ imaging in 3D bioprinted constructs. Advanced Functional Materials 28:1804411.

Waterbury RB, Stanier RY. 1978. Patterns of growth and development in pleurocapsalean cyanobacteria. Microbiological Reviews 42:2–44.

Vousdoukas MI, Velegrakis AF, Plomaritis TA. 2007. Beachrock occurrence, characteristics, formation mechanisms and impacts. Earth Science Reviews 85:23–46.

Zhang Z-C, Li Z-K, Yin Y-C, Li Y, Chen M, Qiu B-S. 2019. Widespread occurrence and unexpected diversity of red-shifted chlorophyll producing cyanobacteria in humid subtropical forest ecosystems. Environmental Microbiology 21:1497–1510.

Zhao C, Gan F, Shen G, Bryant DA. 2015. RfpA, RfpB, and RfpC are the master control elements of far-red light photoacclimation (FaRLiP). Frontiers in Microbiology 6: 1–13.

